# *pbzarr*: cohort-scale storage and analysis of per-base genomic data with Zarr

**DOI:** 10.64898/2026.11.16.767130

**Authors:** Cade Mirchandani, Walker Meyer, Alexis Garretson, Aaron R. Quinlan, Mitchell R. Vollger

## Abstract

Genomic assays now produce per-base signals for cohorts comprising thousands of samples. However, existing genomic data storage formats (e.g., D4, bigWig) were designed primarily for single samples or individual datasets. Consequently, cohort-scale analyses across samples, populations, or species require opening, reading, and merging thousands of files before proceeding to downstream computational analyses. Here, we present *pbzarr*, a format and toolkit that stores per-base genomic data from many samples and assays as a single Zarr store. Across multiple datasets, *pbzarr* simplified cohort construction to a single command and created cohort-scale datasets up to 2.1x faster than existing approaches while simultaneously using 5-6x less memory. More importantly, *pbzarr* compresses signal tracks to only a fraction of the input data size, using only 3.3% of the input size to store per-base coverage for 1,024 short-read genomes. Despite this smaller storage footprint, *pbzarr* maintains fast access to underlying genomic data: region-based queries complete 61x faster than D4 and 6x faster than bigWig, and full-genome scans are as fast or faster than existing formats. Finally, because pbzarr stores are native Zarr arrays, it integrates directly with widely used array libraries (e.g., Xarray and Dask), allowing cohort-scale analyses to be expressed in a single line of Python code using standard array operations instead of a complex, custom pipeline. Together, our results suggest that *pbzarr* is dramatically smaller, substantially simpler, and faster than competing formats for biobank-scale analysis. *pbzarr* is open source and available at https://github.com/pbzarr/pbzarr.

## Introduction

Falling sequencing costs and growing demand for personalized medicine have made population-scale whole-genome sequencing routine. With this increased utilization and access, biobank-scale datasets exceeding 100,000 individuals are increasingly available (Goodwin et al. 2016; Shendure et al. 2017; Byrska-Bishop et al. 2022; All of Us Research Program Genomics Investigators 2024; UK Biobank Whole-Genome Sequencing Consortium 2025). Sequencing has also expanded beyond merely measuring genomic content to study various molecular readouts including gene expression, DNA methylation, and chromatin accessibility across large cohorts and many tissues (Wilks et al. 2021b; GTEx Consortium 2020; ENCODE Project Consortium 2026; Coorens et al. 2025; Somatic Mosaicism across Human Tissues Network et al. 2026).

The scale and dimensionality of current sequencing assays introduce new demands on our methods and genomic formats as cross-sample and cross-assay analysis grows in complexity. A near-universal need across genomic assays is the ability to store signals at every position in the genome, and to date we have used formats like bigWig (Kent et al. 2010), bedGraph, and D4 (Hou et al. 2021) to accomplish this. However, these formats were designed for a single sample, making cross-sample, multimodal, and integrated analysis difficult. With one file per sample, reading a region across many samples means opening, reading, and merging many files. To run in parallel, analyses rely on workflow managers (Mölder et al. 2021; Di Tommaso et al. 2017) or custom scripts that split the genome into pieces and merge the results. General-purpose array formats such as HDF5 (Folk et al. 2011) and Zarr (Miles et al. 2026) offer a natural solution to these problems. They store large N-dimensional arrays as compressed blocks along every axis, so that a program can read, e.g., a particular region in the genome for a subset of samples. These formats are widely used across other scientific fields (Rasp et al. 2024; Moore et al. 2021; Ruan et al. 2024) and are increasingly adopted in genomics. For example, VCF Zarr uses the Zarr format to efficiently read variant calls by variant and by sample (Czech et al. 2025). Zarr is an open, community-governed specification (fiscally sponsored by NumFOCUS, the nonprofit that also supports NumPy, pandas, and SciPy) with independent implementations in Python, Rust, C++, Java, JavaScript, Julia, and R. Data stored in Zarr therefore do not depend on a single software package or vendor, which the FAIR principles require for data to be interoperable and reusable (Wilkinson et al. 2016). Zarr itself, however, defines only how arrays are stored, and no existing specification defines how to represent signal along a genome across many samples and assays as Zarr arrays.

To make analyses across many samples and assays faster and simpler, we developed *pbzarr*, a format and set of tools for base-pair-resolution data as Zarr arrays. *pbzarr* is motivated by the observation that analyses at biobank scale often require indexing across multiple genomic regions, samples, and data types. A *pbzarr* store is composed of tracks, each of which is an array with a genomic axis shared by all tracks and any number of labeled axes such as sample, haplotype, or modification type. We demonstrate that pbzarr stores are smaller than D4 and bigWig files, faster to build and to query across many samples, and substantially simplify analysis. *pbzarr* and its tools support the importing of data from common alignment and signal formats, fast queries by region and by sample, and summaries across samples and assays. Beyond these tools, any Zarr library can read a *pbzarr* store, so improvements to Zarr and its ecosystem apply to *pbzarr* without changes to the format. Together, compact storage, fast queries across samples, and an open, widely supported foundation make *pbzarr* a basis for genomic analyses at biobank scale.

## Results

### Implementation of *pbzarr*

*pbzarr* is a single **A** store for genomic sequencing data that allows for joint computation and storage across data types, genomic position, and samples. For example, a single *pbzarr* store can be used to represent a cohort of individuals jointly for their quantitative signals (e.g., genomic coverage and DNase), while also storing categorical annotations (e.g., positions of repetitive elements) for stratification in data analysis (**Fig. 1A**). Importantly, this allows for arbitrary computation on or across *tracks* and their shared dimensions. For example, an intersection of DNase values across a cohort with repetitive element annotations is a single matrix operation resulting in DNase values stratified by repetitive elements (**Fig. 1B)**. *pbzarr* consists of four parts: a format specification, a Rust library (crate: *pbzarr*), a Python package (PyPI: *pbzarr*), and a command-line tool (*pbz*).

**Figure 1.**
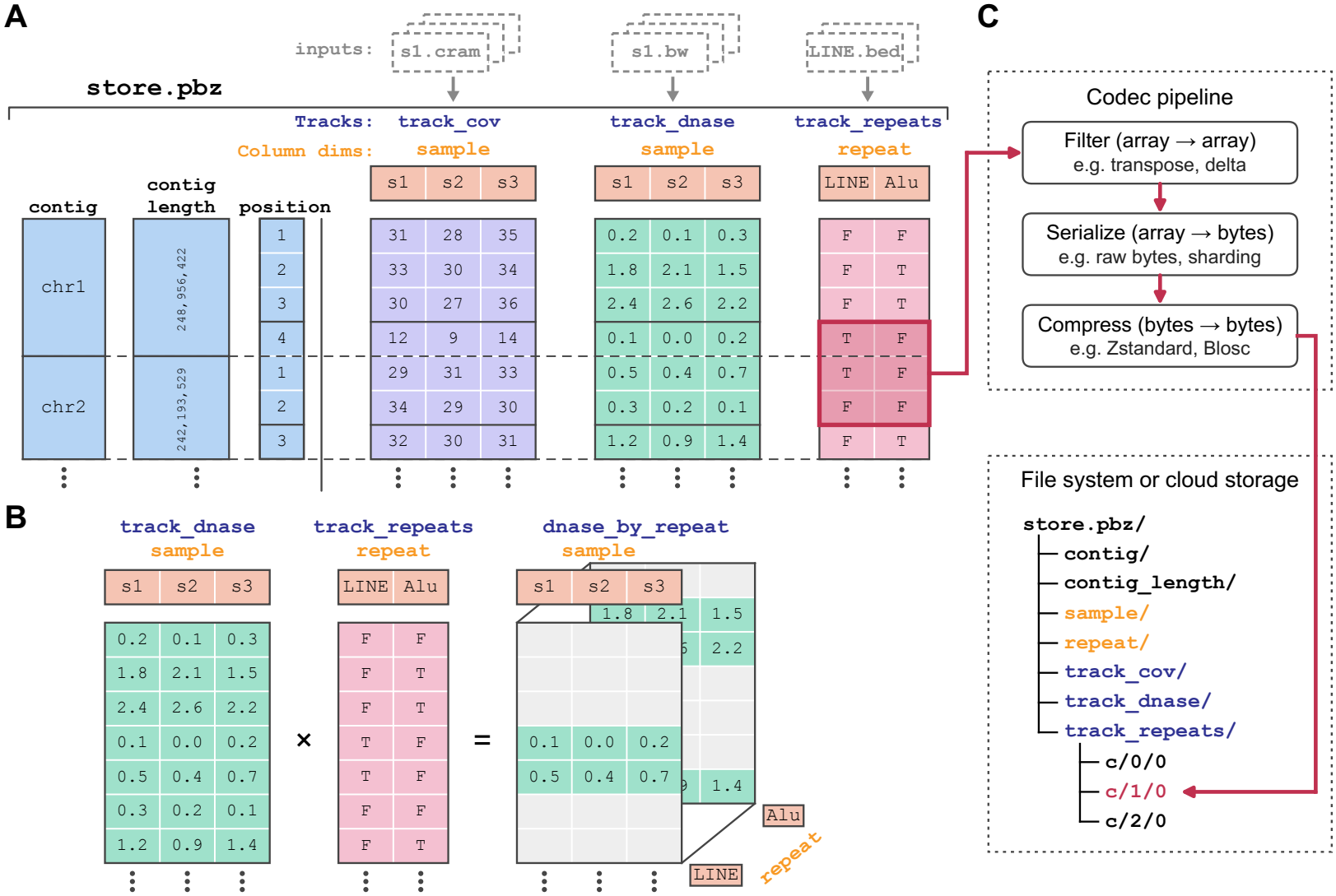
The *pbzarr* format. **(A)** A pbzarr store holds a reference genome, defined by contig IDs and contig lengths (blue), and any number of tracks. Tracks share the position axis, which runs over every base in reference order (dashed line, contig boundary), and can hold quantitative signals (purple, green) or categorical annotations (red) along a labeled second dimension. **(B)** Intersecting the DNase track with the repeat track gives DNase values for each sample stratified by repeat class. **(C)** Each chunk of a track (red outline in A) passes through a Zarr codec pipeline and is stored as one object named by its chunk index (e.g., c/1/0), on a file system or in cloud object storage.

#### Format

Formally, a *pbzarr* store is a Zarr v3 group that holds a reference genome defined by contig IDs and contig lengths, and track arrays (*tracks*) with a position dimension (*position*), followed by optional labeled dimensions (*column dim*) such as sample (**Fig. 1A**, https://pbzarr.github.io/reference/format/). Each track is a Zarr array, which is split into blocks of consecutive positions and samples (chunks) that are compressed and stored independently (**Fig. 1C**). By default, a chunk holds 65,536 positions of one sample, and many chunks are grouped into one file (a shard), which keeps the number of files small while a program can still read a single chunk. Because a *pbzarr* store is an ordinary Zarr store, it can be a directory on a file system, a ZIP file, or a location in cloud object storage, and any Zarr v3 library can read it without *pbzarr* code. By convention, the name of a *pbzarr* store ends in “.pbz.”

#### Rust library

The Rust library (https://crates.io/crates/pbzarr) uses zarrs (Deakin 2026) to create, read, and do analysis on *pbzarr* stores. The Rust library can be used to directly build stores from many common bioinformatics formats: BAM, CRAM, D4, bigWig, and tabix-indexed BED files. For alignments, it calculates per-base depth directly from the reads, and can also count the bases, insertions, and deletions observed at each position. All input files are read in one multi-threaded process that holds only a bounded number of genomic segments in memory at once, so memory use depends on the number of threads and changes little with the number of samples.

#### Python package

The Python package (https://pypi.org/project/pbzarr/) supports analyses that are more complex than the summaries the command-line tool provides. In Python, a store opens as a single Xarray Dataset (Hoyer and Hamman 2017), in which each track is a lazy, Dask-backed array (Rocklin 2015) with named dimensions and sample labels. Standard array operations therefore apply to tracks without *pbzarr*-specific code and use Dask to run in parallel. By design, the Python package only adds operations that need genomic coordinates: selecting a region by contig and position, summarizing a track over many regions, and writing a result back to the store as a new track. Developers can also use the Python package (or Rust library) to read and write *pbzarr* stores from their own tools, so that existing software can adopt the format without implementing the specification itself.

#### Command-line

The command-line tool, *pbz,* exposes aspects of the Rust library through a set of subcommands. *pbz import* builds a store from any number of input files, and *pbz append* adds samples to an existing store. *pbz stat* calculates the mean, minimum, maximum, median, or histogram of a track for each sample, over whole contigs or the regions of a BED file. *pbz view* writes a track as bedGraph-like text.

### Benchmarking creation of cohort-scale datasets with *pbzarr*

To benchmark the creation and storage of large datasets, we compared ***pbzarr*** with D4 (Hou et al. 2021) and bigWig (Kent et al. 2010). We benchmarked against these tools because D4 is the most recent format designed for dense depth data, and bigWig is the most widely used format for per-base signal. We evaluated the formats and their respective tools using three datasets: 1,024 high-coverage short-read genomes from the 1000 Genomes Project (Byrska-Bishop et al. 2022), 1,019 long-read genomes from the same samples (Schloissnig et al. 2025), and 60 long-read RNA-seq samples (Réal et al. 2026).

#### Faster and simpler dataset creation

The bigWig and D4 workflows first compute depth for one sample at a time using megadepth (Wilks et al. 2021a) and mosdepth (Pedersen and Quinlan 2018), respectively, and write one file per sample (**Fig. 2A**). D4 then merges these files into a cohort file with d4tools, whereas bigWig has no cohort file and remains 1,024 separate files. *pbzarr* replaces these steps with one command, *pbz import*, which reads all alignments and writes the cohort store directly, with no per-sample intermediates. For 1,024 short-read genomes on 256 cores, building the pbzarr store took 59 min, compared with 91 min for bigWig and 121 min for D4 (1.5- and 2.1-fold faster; **Fig. 2B**). *pbzarr* was also the fastest to build for long-read genomes (1.2-fold faster than both bigWig and D4), but the slowest for long-read RNA (4.7- and 1.3-fold slower than bigWig and D4; **Fig. S1**). Peak memory was 177 GB for *pbzarr*, compared with 924 GB for bigWig and 1,068 GB for D4 (5.2- and 6.0-fold lower). The D4 and bigWig tools run one process per sample, so their memory grew with the number of concurrent jobs and plateaued only once all 256 cores were in use. In contrast, *pbz import* memory grew only modestly with cohort size (**Fig. 2C**)

#### Improved compression

*pbzarr* stores were substantially smaller than D4 and bigWig, and this size improvement grew with the number of samples (**Fig. 2D**). For 1,024 short-read genomes (16.6 TB of CRAM input), the ***pbzarr*** store took only 0.54 TB, 3.3% of the input, compared with 2.9 TB (18%) for D4 and 4.4 TB (26%) for bigWig. Furthermore, we demonstrated that this advantage is not unique to whole-genome short-read datasets by extending this analysis to whole-genome and RNA long-read cohorts (**Fig. 2E**). For 1,019 long-read genomes (6.1 TB of input), *pbzarr* took 9.5% of the input, compared with 46% for D4 and 58% for bigWig. For 60 long-read RNA-seq samples (131 GB of input), it took 1.2%, compared with 125% for D4 (**Supplemental note 1**) and 2.7% for bigWig. This difference comes primarily from how values are encoded. D4 stores each base in a fixed number of bits, whereas pbzarr compresses each chunk with a numerical compressor, pcodec (Loncaric et al. 2025), which finds the long runs of equal depth along the genome

Importantly, for D4 and bigWig to compete with *pbzarr* in speed, as in this analysis, we had to develop a pipeline for multi-sample cohort creation, whereas *pbzarr* created each dataset with a single command.

**Figure 2.**
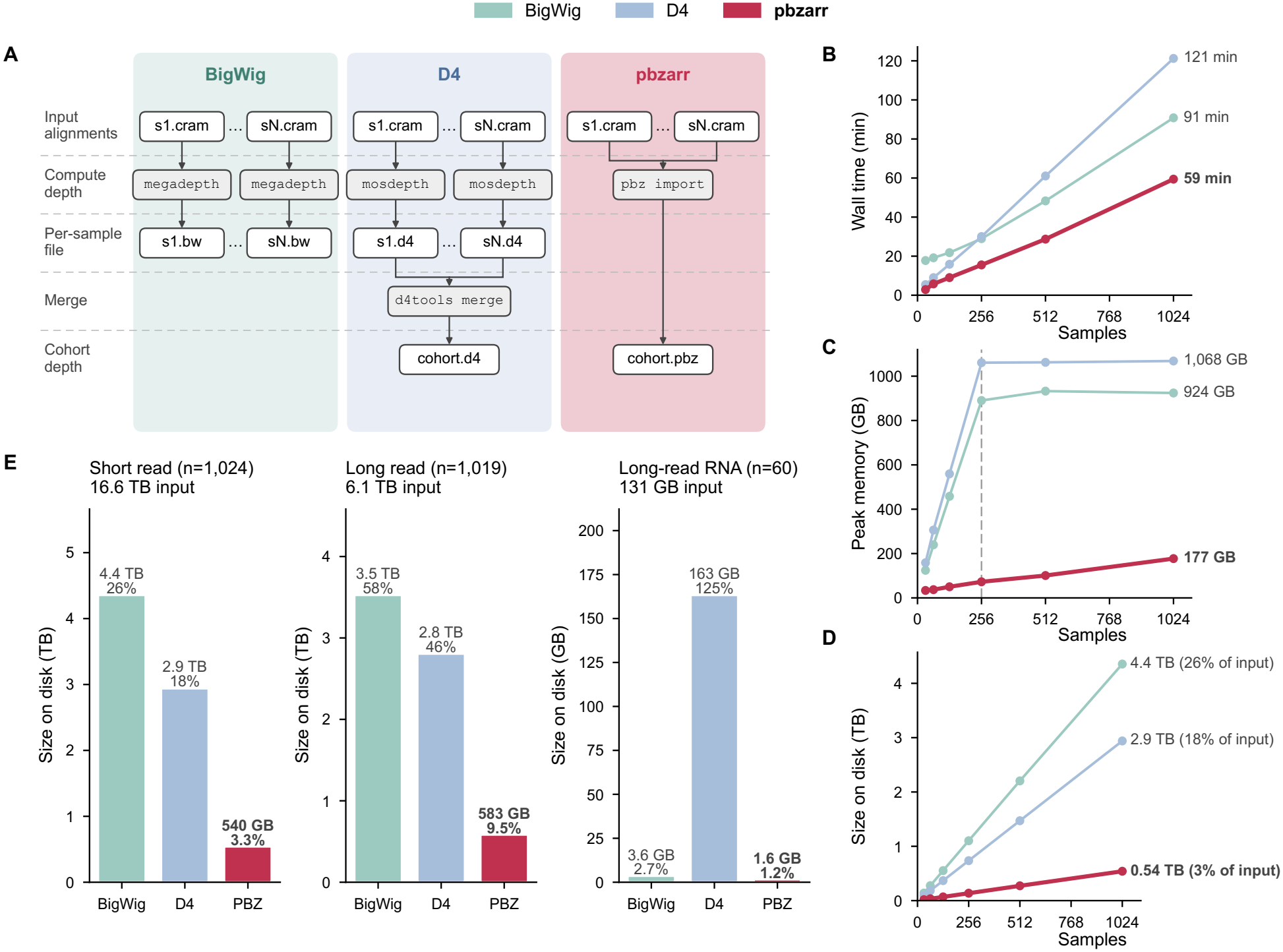
Cohort-scale depth storage with pbzarr. **(A)** Workflows for computing per-base depth across a cohort. **(B, C, D)** Wall time, peak memory, and size on disk, as a function of the number of 1000 Genomes Project (1KG) short-read samples. The dashed line marks the sample count at which all 256 cores are in use. **(E)** Size on disk for three 1KG cohorts: short-read whole-genome sequencing (n=1,024), ONT long-read whole-genome sequencing (n=1,019), and ONT long-read RNA sequencing (n=60). Labels give the absolute size and the size as a percentage of the input alignments.

### Benchmarking full scans and indexed reads of *pbzarr* stores

To demonstrate that *pbzarr* does not sacrifice access speed for compression, we benchmarked computing the average depth of the entire genome across multiple genomes (full scan) and calculating per-region averages (region-scan) (**Fig. 3A**). In this analysis, we compared *pbzarr* to D4 (d4tools) and bigWig (pyBigWig) on calculating mean coverage for 1,024 short-read genomes both across the entire genome and in 10,000 random windows of 10 kbp each. We did not use the precomputed summaries that each format can store, so that each scan measured the cost of reading the per-base values.

**Figure 3.**
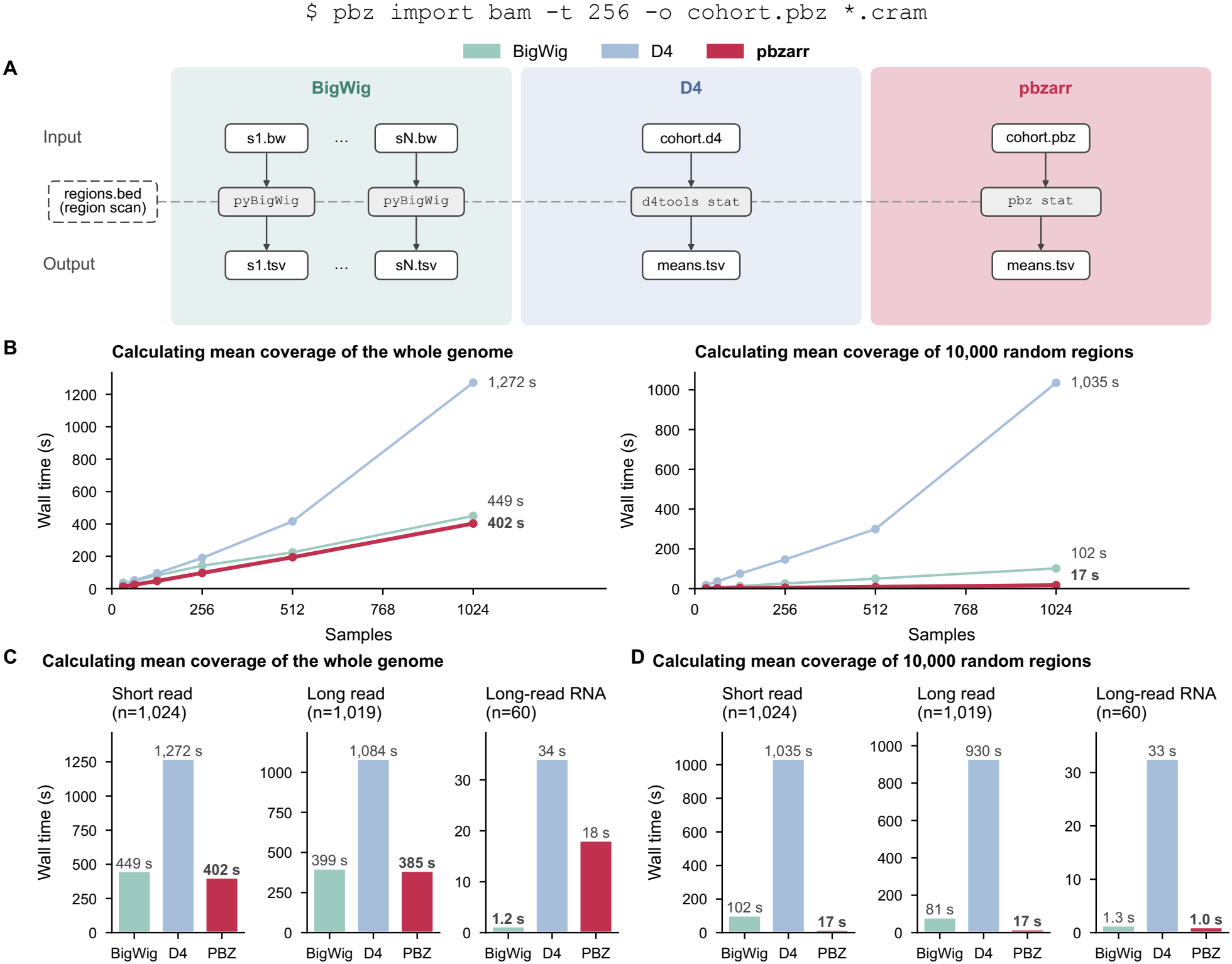
Computing mean depth across a cohort. **(A)** Workflows for computing mean depth per sample across a cohort. **(B)** Wall time as a function of the number of 1000 Genomes Project (1KG) short-read samples, for the whole genome (left) and for 10,000 random regions (right). **(C, D)** Wall time for the whole genome (C) and for 10,000 random regions (D) in three 1KG cohorts: short-read whole-genome sequencing (n=1,024), ONT long-read whole-genome sequencing (n=1,019), and ONT long-read RNA sequencing (n=60). Bold labels mark the fastest tool. `$ pbz import bam -t 256 -o cohort.pbz *.cram`

#### Full scans

For a full scan of 1,024 short-read genomes, *pbzarr* took 402 s, compared with 449 s for bigWig and 1,272 s for D4 (**Fig. 3B**). *pbzarr* was therefore moderately faster than bigWig and substantially faster than D4, though D4 does appear to lose much of its read performance in the merged D4 format compared to individual D4 files (**Figs. S2 and S3**). As previously emphasized, this moderate increase in speed comes with an 8.1- and 5.4-fold smaller *pbzarr* store (**Fig. 2D**). The long-read genomes gave the same ranking (385 s for *pbzarr*, 399 s for bigWig, and 1,084 s for D4; **Fig. 3C**), and the long-read RNA cohort was the exception where bigWig took 1.2 s, compared with 18 s for *pbzarr* and 34 s for D4. This is likely because RNA-seq depth is zero across most of the genome, and bigWig stores only the intervals with signal, whereas *pbzarr* and D4 store a value for every base.

#### Region scans

The random 10,000 windows cover 98.5 Mbp, or about 3% of the genome, so formats with efficient random access should finish in a small fraction of their full-scan time. For 1,024 short-read genomes, *pbzarr* took 17 s, 24-fold less than its full scan (**Fig. 3B, C**). This was 6.0-fold faster than bigWig (102 s) and 61-fold faster than D4 (1,035 s), whose region scan still took 81% of its full-scan time. The long-read genomes behaved similarly (17 s for *pbzarr*, 81 s for bigWig, and 930 s for D4). For long-read RNA, *pbzarr* and bigWig were comparable (1.0 s and 1.3 s), and D4 took 33 s.

Therefore, the smaller *pbzarr* stores do not come at the cost of read speed, and across the three cohorts, *pbzarr* was the fastest format for region queries and full scans of whole-genome data.

### Use case: a cohort-level false discovery rate for chromatin accessibility with *pbzarr*

To demonstrate an analysis that is difficult with per-sample files, we used *pbzarr* to calculate a single false discovery rate (FDR) for regulatory elements called across nine Fiber-seq samples. Fiber-seq marks accessible DNA on single chromatin fibers (Stergachis et al. 2020), and FIRE (Vollger et al. 2026) combines these fibers into a per-base accessibility score for each sample. To set a threshold, FIRE shuffles the fibers along the genome, rescores them, and defines the FDR at each score as the number of shuffled bases at or above that score divided by the number of real bases. However, thresholds set separately for each sample do not control the FDR of a combined call set, because the union of per-sample peaks is more likely to overlap in real regulatory elements than false positives.

We therefore stored the real and shuffled scores of all nine samples as two tracks in one *pbzarr* store, took the maximum of each across samples at every base, and recalculated the FDR on the merged tracks (**Fig. 4**). This recalibrated threshold improved overlap with known ENCODE cis-regulatory elements for regions with FDR ≤ 5% (92.1% compared with 86.1% for the union), consistent with improved specificity. Notably, this task took less than 3 minutes and required only a few lines of Python code, despite the dataset comprising 55.8 billion values (**Box 1 and Supplemental note**).

**Box 1.**

pbzarr code for calculating FDR.

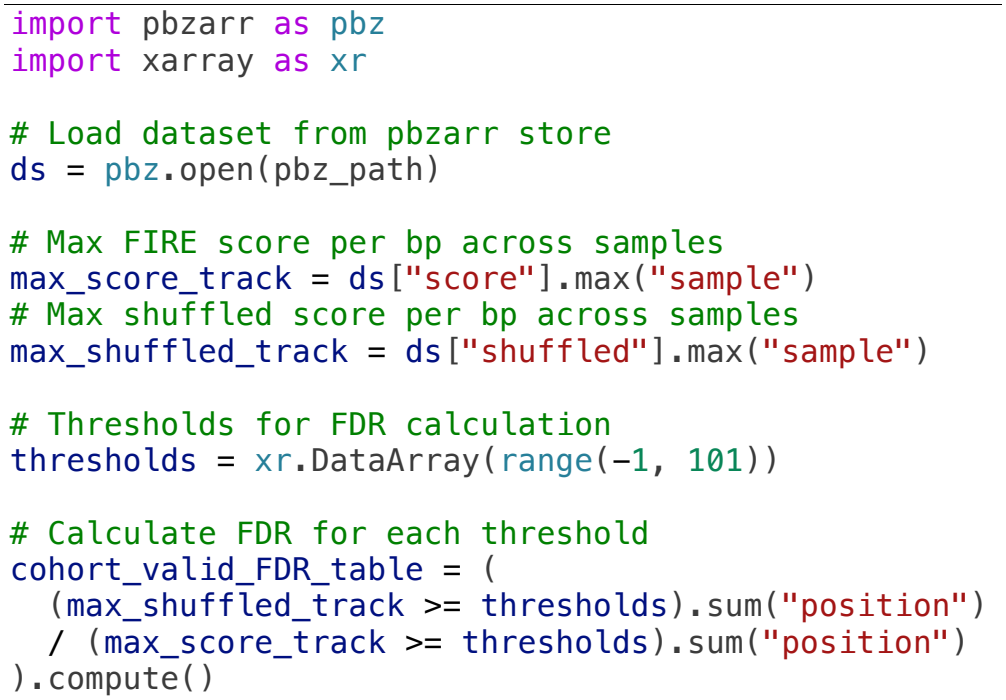

**Figure 4.**
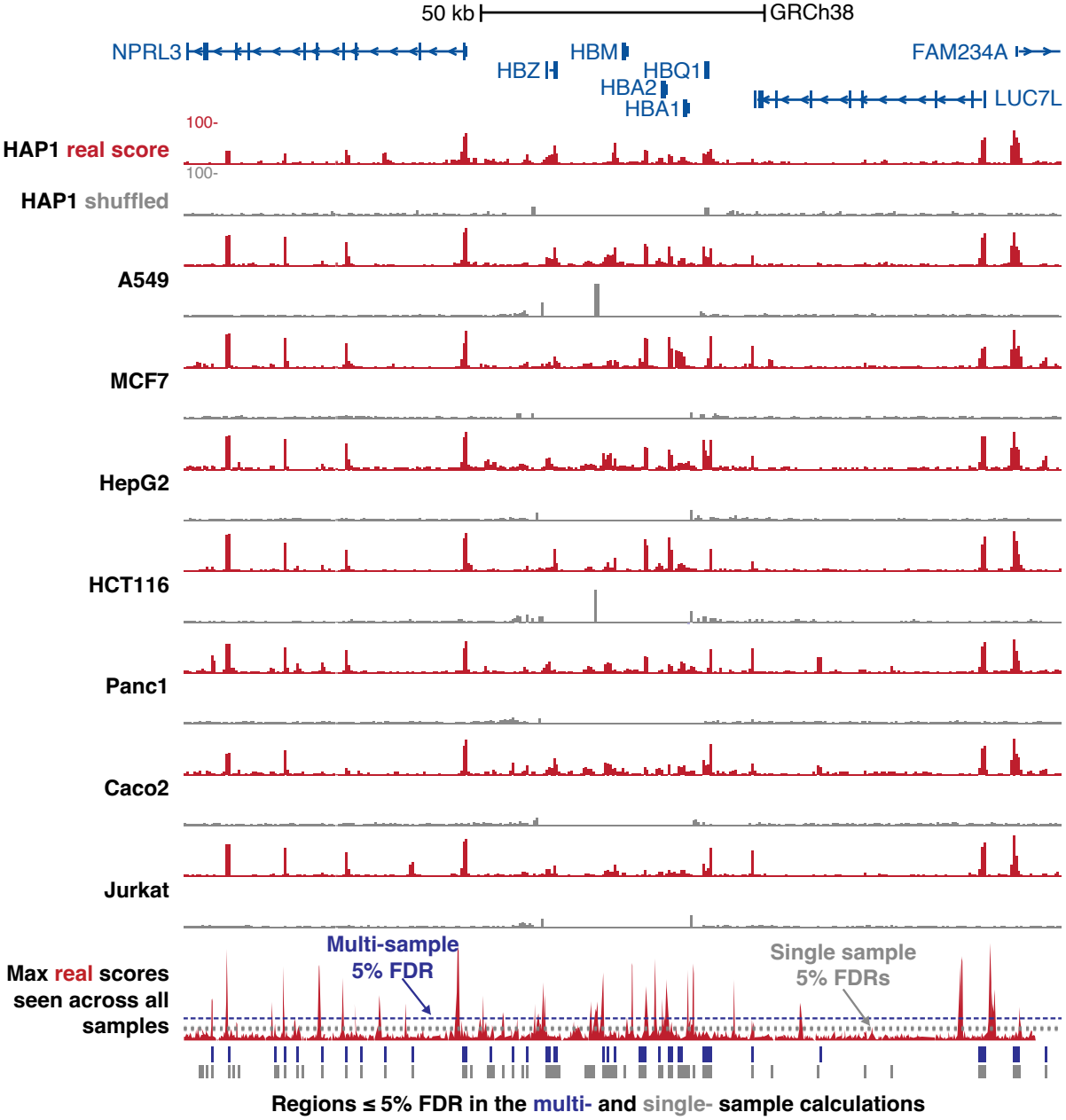
A unified false discovery rate across samples with pbzarr. Fiber-seq FIRE scores (red) and scores from shuffled reads (gray) at the α-globin locus (GRCh38) for a subset of cell lines. The bottom track shows the maximum score across all samples with the 5% FDR thresholds estimated from each sample alone (gray dashed lines) and from the merged maximum tracks (blue dashed line). Bars below mark the regions that pass the multi-sample (blue) and single-sample (gray) thresholds.

## Discussion

Here, we developed the command-line tool, libraries, and specification for *pbzarr,* a format for storing and computing on per-base genomic data at biobank scale. *pbzarr* is competitive and, in many cases, outperforms other file formats for read and write speed across multiple use cases. However, beyond these baseline improvements, *pbzarr* makes three conceptual advances worth highlighting:

1. **Multi-sample and multimodal.** *pbzarr’s* design lets it organize data in one store across samples, data types, and/or annotations. We demonstrate that this allows for easy genomic stratification of results across many samples (**Fig. 1B**) or jointly calculating statistics across multiple samples (**Fig. 4**). Furthermore, *pbzarr* stores can be appended to, so additional samples added in the future to a large cohort don’t require costly re-computation on the whole dataset.
2. **Storage flexibility.** *pbzarr* across benchmarks is substantially smaller than the other tested formats (2– 8-fold smaller), and importantly, in the case of storing whole-genome coverage profiles, *pbzarr* is only ∼3% of the size of the input data (CRAMs), making it practical to save coverage profiles rather than constantly recomputing them. To demonstrate this, and the cloud capabilities of *pbzarr,* we now host coverage profiles for all 3,202 high-coverage samples from the 1000 Genomes Project in a small S3 store for public use (https://pbzarr.github.io/get-started/1000-genomes/).
3. **Simplicity.** *pbzarr* enables a cohort-scale analysis of, e.g., coverage in a line of Python rather than a pipeline. This is possible because of three implementation details of *pbzarr*. One, Zarr (and *pbzarr*) stores are chunked, so there is a natural and predefined unit of work. This is in contrast to many genomic analyses, where careful effort is needed to divide computation across datasets without sacrificing efficiency. Two, *pbzarr* is indexed both along the genome and along its labeled dimensions so that computation can be effectively limited to either a subset of the genome, a subset of samples, or both. Finally, *pbzarr* parallelizes almost every task with no additional user effort, thanks to its shared infrastructure with Zarr (Miles et al. 2026), Xarray (Hoyer and Hamman 2017), Dask (Rocklin 2015), and others.

### Limitations and future directions of *pbzarr*

*pbzarr* does face several important limitations. First, for best performance, *pbzarr* needs to remain a multi-file store. The Zarr community is already working to address this using a zipped format, but it currently has performance limitations. Second, *pbzarr* is not well suited for interval or per-molecule data at this time, and any such data would have to be converted to per-base signal before use with *pbzarr*. Finally, *pbzarr* does not yet support sparse representations, and this can make it slower than interval-based formats (bigWig) on sparse datasets.

However, *pbzarr* also offers many advantages not fully explored here. For example, GPUs can read *pbzarr* directly, complementing large-scale machine learning in genomics, and Zarr’s cloud-native capabilities allow hosting genomic resources that can be directly computed on, from anywhere (e.g., https://pbzarr.github.io/get-started/1000-genomes/). Finally, *pbzarr* will directly benefit from future advances in Zarr and the surrounding ecosystem. Overall, *pbzarr* is a fast, extremely storage-efficient format for per-base genomic data that enables simple analysis of large and complex cohorts.

## Methods

### Benchmarks

In this section, we describe the methodology used for the benchmarks in Figures 2 and 3. The benchmarks use three public datasets, chosen to cover different sequencing technologies and depth profiles: dense depth from short-read and long-read genomes, and sparse depth from long-read RNA sequencing, where most of the genome has no coverage. For short-read benchmarking, we used the CRAM files from the 1000 Genomes 30x on GRCh38 data collection (Byrska-Bishop et al. 2022). To measure how build time, memory, output size, and scan time change with the number of samples, we built cohorts of 32, 64, 128, 256, 512, and 1,024 samples from this dataset. For long reads, we used 1,019 Oxford Nanopore genomes from the 1KG_ONT_VIENNA data collection (Schloissnig et al. 2025), and we used 60 Oxford Nanopore direct RNA samples (Réal et al. 2026). For both the whole-genome long-read and long-read RNA datasets, we only performed benchmarking on all samples in each dataset.

We ran all benchmarks on a dedicated server with two AMD EPYC 9755 processors (256 cores in total) and 1.5 TB of RAM running Rocky Linux 8.10. For each cohort, we built each format’s dataset in turn on the server’s local disk, queried it, and deleted it before building the next. For *pbzarr*, we used a single *pbz import v0.6.0* command to read all the cohort’s alignment files into one store. We used *mosdepth v0.3.14* (Pedersen and Quinlan 2018) to generate per-sample D4 depth files, and merged them into one cohort file using *d4tools merge v0.3.11* (Hou et al. 2021). For bigWig, we used *megadepth v1.2.0* (Wilks et al. 2021a) to create per-sample bigWigs, which together make up the bigWig dataset. *pbz import* used all 256 cores in one process, and we used *Snakemake v9.27.0* (Mölder et al. 2021) to parallelize the *mosdepth* and *megadepth* jobs, dividing the cores evenly among each tool’s jobs.

We then queried each dataset in two ways: a full scan, computing each sample’s mean depth over the whole genome, and an indexed read, computing each sample’s mean depth over 10,000 random 10-kb regions generated with *BEDtools v2.31.1* (Quinlan and Hall 2010). For *pbzarr*, we used *pbz stat*; for D4, we used *d4tools stat* with the option *--no-index* for the full scan to ensure we measured the time to read the whole file. Finally, for bigWig, we used a custom Python script with *pyBigWig*, using the *exact* option to avoid using bigWig’s built-in zoom summaries. Each query job was run with 256 cores.

To prevent data cached by earlier runs from affecting the measurements, we evicted each run’s input files from the operating system’s page cache with posix_fadvise (POSIX_FADV_DONTNEED) before every build and query. For each build and query, we measured elapsed wall-clock time (including flushing output to disk), total CPU time (the sum of the user and system times of all processes) and peak memory (the resident set size summed over all running processes). For each build, we also measured the dataset size on disk. We ran the full benchmark three times for each cohort and ran each query three times within each run, and we report medians.

### Cohort-level FDR for Fiber-seq

We obtained Fiber-seq data for nine human cell lines (A549, MCF-7, HepG2, HCT116, Caco-2, Panc1, Jurkat, HAP1, and K562) from the Fiber-seq open-access repository (https://s3.kopah.uw.edu/fiberseq-open-access/index.html) and processed them with FIRE on GRCh38. For each sample, FIRE produces a per-base score and a null score, which it computes after moving every read to a random position on the same chromosome, excluding regions of unreliable coverage. We imported the real and null scores of all samples into one *pbzarr* store as two tracks that share a sample dimension. For a score threshold *t*, we defined the FDR as the number of bases with a null score of at least *t* divided by the number of bases with a real score of at least *t*, and we used the smallest *t* with an FDR of 5% or less (Vollger et al. 2026).

For per-sample thresholds, we applied this calculation to each sample individually, and the union call set retained a base when any sample met its threshold. For the recalibrated threshold, we took the maximum real score and maximum null score across the nine samples at each base, then repeated the calculation on these merged tracks. To measure overlap with regulatory annotations, we intersected each call set with the ENCODE registry of candidate cis-regulatory elements using BEDtools (Quinlan and Hall 2010) and reported the fraction that overlapped an ENCODE element.

## Conflict of interest

None declared.

## Acknowledgements

We thank Andrew B. Stergachis, Jane Ranchalis, and Shane Neph for generating and publicly sharing Fiber-seq on ENCODE cell lines, and Google for funding the generation of these datasets. We thank the 1000 Genomes Project for generating the benchmarking datasets, and Per Unneberg and Chris Condon for feedback on the development and early implementation of *pbzarr*.

## Funding

This work was supported by the National Institute of General Medical Sciences (NIGMS) via a Pathway to Independence Award to M.R.V. (R00GM155552). The computational resources used were partially funded by the NIH Shared Instrumentation Grants S10OD034321 and S10OD021644.

## Software availability

Documentation for *pbzarr* can be found at https://pbzarr.github.io/. *pbzarr* is open source under the MIT license at https://github.com/pbzarr/pbzarr, which also hosts the specification (https://github.com/pbzarr/pbzarr/tree/main/spec). It provides a Rust library (https://crates.io/crates/pbzarr), a Python package (pip install pbzarr; https://pypi.org/project/pbzarr), and the *pbz* command-line tool, available as prebuilt binaries on GitHub. Benchmarking code can be found at https://github.com/pbzarr/pbzarr-benchmarking.

## Notes

https://pbzarr.github.io/

### Competing Interest Statement

The authors have declared no competing interest.

## References

All of Us Research Program Genomics Investigators. 2024. Genomic data in the All of Us Research Program. Nature 627: 340–346. 10.1038/s41586-023-06957-x.

Byrska-Bishop M, Evani US, Zhao X, Basile AO, Abel HJ, Regier AA, Corvelo A, Clarke WE, Musunuri R, Nagulapalli K, et al. 2022. High-coverage whole-genome sequencing of the expanded 1000 Genomes Project cohort including 602 trios. Cell 185: 3426–3440.e19. 10.1016/j.cell.2022.08.004.

Coorens THH, Oh JW, Choi YA, Lim NS, Zhao B, Voshall A, Abyzov A, Antonacci-Fulton L, Aparicio S, Ardlie KG, et al. 2025. The Somatic Mosaicism across Human Tissues Network. Nature 643: 47–59. 10.1038/s41586-025-09096-7.

Czech E, Tyler W, White T, Jeffery B, Millar TR, Elsworth B, Guez J, Hancox J, Karczewski KJ, Miles A, et al. 2025. Analysis-ready VCF at Biobank scale using Zarr. Gigascience 14: giaf049. 10.1093/gigascience/giaf049.

Deakin L. 2026. Zarrs: A high-performance Rust library for the Zarr array storage format. Zenodo 10.5281/ZENODO.12719776.

Di Tommaso P, Chatzou M, Floden EW, Barja PP, Palumbo E, Notredame C. 2017. Nextflow enables reproducible computational workflows. Nat Biotechnol 35: 316–319. 10.1038/nbt.3820.

ENCODE Project Consortium. 2026. The Encyclopedia of DNA elements. bioRxivorg. 10.64898/2026.07.06.731365.

Folk M, Heber G, Koziol Q, Pourmal E, Robinson D. 2011. An overview of the HDF5 technology suite and its applications. In Proceedings of the EDBT/ICDT 2011 Workshop on Array Databases, ACM, New York, NY, USA 10.1145/1966895.1966900.

Goodwin S, McPherson JD, McCombie WR. 2016. Coming of age: ten years of next-generation sequencing technologies. Nat Rev Genet 17: 333–351. 10.1038/nrg.2016.49.

GTEx Consortium. 2020. The GTEx Consortium atlas of genetic regulatory effects across human tissues. Science 369: 1318–1330. 10.1126/science.aaz1776.

Hou H, Pedersen B, Quinlan A. 2021. Balancing efficient analysis and storage of quantitative genomics data with the D4 format and d4tools. Nat Comput Sci 1: 441–447. 10.1038/s43588-021-00085-0.

Hoyer S, Hamman J. 2017. xarray: N-D labeled Arrays and Datasets in Python. J Open Res Softw 5: 10. 10.5334/jors.148.

Kent WJ, Zweig AS, Barber G, Hinrichs AS, Karolchik D. 2010. BigWig and BigBed: enabling browsing of large distributed datasets. Bioinformatics 26: 2204–2207. 10.1093/bioinformatics/btq351.

Loncaric M, Jeppesen N, Zinberg B. 2025. Pcodec: Better compression for numerical sequences. arXiv [csIT]. 10.48550/arXiv.2502.06112.

Miles A, Bennett D, jakirkham, Stansby D, Orfanos DP, Hamman J, Bussonnier M, Moore J, Jones M, Augspurger T, et al. 2026. zarr-developers/zarr-python: v3.4.2. Zenodo 10.5281/ZENODO.23695418.

Mölder F, Jablonski KP, Letcher B, Hall MB, Tomkins-Tinch CH, Sochat V, Forster J, Lee S, Twardziok SO, Kanitz A, et al. 2021. Sustainable data analysis with Snakemake. F1000Res 10: 33. 10.12688/f1000research.29032.2.

Moore J, Allan C, Besson S, Burel J-M, Diel E, Gault D, Kozlowski K, Lindner D, Linkert M, Manz T, et al. 2021. OME-NGFF: a next-generation file format for expanding bioimaging data-access strategies. Nat Methods 18: 1496–1498. 10.1038/s41592-021-01326-w.

Pedersen BS, Quinlan AR. 2018. Mosdepth: quick coverage calculation for genomes and exomes. Bioinformatics 34: 867–868. 10.1093/bioinformatics/btx699.

Quinlan AR, Hall IM. 2010. BEDTools: a flexible suite of utilities for comparing genomic features. Bioinformatics 26: 841–842. 10.1093/bioinformatics/btq033.

Rasp S, Hoyer S, Merose A, Langmore I, Battaglia P, Russell T, Sanchez-Gonzalez A, Yang V, Carver R, Agrawal S, et al. 2024. WeatherBench 2: A benchmark for the next generation of data-driven global weather models. J Adv Model Earth Syst 16. 10.1029/2023ms004019.

Réal A, Brown A, Yung GP, Borel C, Lykoskoufis N, Seebach J, Dermitzakis E, Ramisch A, Viñuela A. 2026. Direct long-read RNA sequencing uncovers functional variation affecting transcript production and RNA modifications. Research Square. 10.21203/rs.3.rs-4613444/v2.

Rocklin M. 2015. Dask: Parallel Computation with Blocked algorithms and Task Scheduling. In Proceedings of the Python in Science Conference, pp. 126–132, SciPy https://proceedings.scipy.org/articles/Majora-7b98e3ed-013.pdf (Accessed November 12, 2026).

Ruan X, Mueller M, Liu G, Görlitz F, Fu T-M, Milkie DE, Lillvis JL, Kuhn A, Gan Chong J, Hong JL, et al. 2024. Image processing tools for petabyte-scale light sheet microscopy data. Nat Methods 21: 2342–2352. 10.1038/s41592-024-02475-4.

Schloissnig S, Pani S, Ebler J, Hain C, Tsapalou V, Söylev A, Hüther P, Ashraf H, Prodanov T, Asparuhova M, et al. 2025. Structural variation in 1,019 diverse humans based on long-read sequencing. Nature 644: 442–452. https://www.nature.com/articles/s41586-025-09290-7 (Accessed November 12, 2026).

Shendure J, Balasubramanian S, Church GM, Gilbert W, Rogers J, Schloss JA, Waterston RH. 2017. DNA sequencing at 40: past, present and future. Nature 550: 345–353. 10.1038/nature24286.

Somatic Mosaicism across Human Tissues Network, Sedlazeck FJ, Coorens THH, Park PJ, Stergachis AB. 2026. Integrated map of somatic mosaicism across human tissues in 25 individuals. bioRxivorg 2026.09.01.748636. 10.64898/2026.09.01.748636v1.

Stergachis AB, Debo BM, Haugen E, Churchman LS, Stamatoyannopoulos JA. 2020. Single-molecule regulatory architectures captured by chromatin fiber sequencing. Science 368: 1449–1454. 10.1126/science.aaz1646.

UK Biobank Whole-Genome Sequencing Consortium. 2025. Whole-genome sequencing of 490,640 UK Biobank participants. Nature 645: 692–701. 10.1038/s41586-025-09272-9.

Vollger MR, Swanson EG, Neph SJ, Mallory BJ, Dubocanin D, Ho C-H, Cheng Y-HH, Ranchalis J, Munson KM, Sedeno-Cortes AE, et al. 2026. Somatic epimutations cap genetic determinism in the human diploid chromatin epigenome. Cell 189: 6613–6629.e9. 10.1016/j.cell.2026.09.034 (Accessed October 15, 2026).

Wilkinson MD, Dumontier M, Aalbersberg IJJ, Appleton G, Axton M, Baak A, Blomberg N, Boiten J-W, da Silva Santos LB, Bourne PE, et al. 2016. The FAIR Guiding Principles for scientific data management and stewardship. Sci Data 3: 160018. 10.1038/sdata.2016.18.

Wilks C, Ahmed O, Baker DN, Zhang D, Collado-Torres L, Langmead B. 2021a. Megadepth: efficient coverage quantification for BigWigs and BAMs. Bioinformatics 37: 3014–3016. 10.1093/bioinformatics/btab152.

Wilks C, Zheng SC, Chen FY, Charles R, Solomon B, Ling JP, Imada EL, Zhang D, Joseph L, Leek JT, et al. 2021b. recount3: summaries and queries for large-scale RNA-seq expression and splicing. Genome Biol 22: 323. 10.1186/s13059-021-02533-6.

