## Supplemental Materials for "*pbzarr*: cohort-scale storage and analysis of per-base genomic data with Zarr"

### Supplemental notes

#### 1. D4 file size for sparse data.

D4 stores depth in a primary table with a fixed number of bits for every base in the genome. Values that do not fit in that width go to a secondary table. d4tools offers a sparse mode (d4tools create --sparse, equivalent to -z -R0-1). It makes the primary table one bit wide and compresses the secondary table, which suits data in which most of the genome has no coverage, such as RNA-seq. mosdepth writes D4 files with its --d4 option but does not offer this mode.

For the long-read RNA dataset, the D4 files we built with mosdepth were therefore larger than the input alignments. For one sample (HG00260), the mosdepth D4 file was 2.72 GB, 124% of the 2.2 GB BAM, or about 7 bits per base. The same sample written with d4tools create --sparse was 4.3 MB (0.2%). The 60-sample D4 cohort, merged from mosdepth files, was 163 GB (125% of the input).

We used mosdepth rather than d4tools create to build all D4 datasets so that every format stored the same depths. With the same read filters, mosdepth, megadepth, and *pbz import* agreed at more than 99.9% of covered bases. d4tools differed at 27% of covered bases for a short-read sample and 96% for an RNA sample, because it does not correct for overlapping read mates; it counts deletions and, for RNA, spliced-out introns as covered.

#### 2. FDR calculation with a multi-sample Fiber-seq *pbzarr*.

When merging per-base datasets for cohort-level analysis, false discovery rates (FDR) from individual samples compound upon merging. It is therefore useful to recompute FDRs in a single merged per-base dataset rather than naively referencing individual samples' FDR scores, as they are no longer valid. This is the case because when call sets from multiple samples are merged, true positives are likely to overlap between samples, whereas false positives are unlikely to overlap. Thus, the merged call set will have a higher FDR than any of the individual samples.

We executed a cohort-level FDR calculation using *pbzarr* with nine Fiber-seq samples, computing 101 FDR scores across a 223 GB uncompressed dataset. This computation took 2 minutes and 55 seconds using 416 threads. Parallel computation of these 101 scores across 2 data tracks, 3.1 Gbp, and 9 samples (55.8 billion values) is prohibitively difficult with single-sample approaches, but becomes easy with pbzarr.

To test whether recalculating FDR on the merged data is useful, we compared overlap with the ENCODE Registry of candidate cis-regulatory elements (cCREs) between intervals that pass the recalculated FDR and a naive merge of intervals that pass the per-sample FDR. The cCREs are regions that ENCODE identified as likely regulatory elements, so an interval that overlaps a cCRE is less likely to be a false positive. Each FDR-passing BED interval (FDR <= 0.5) in these two approaches was classified as cCRE-associated if it overlapped at least one cCRE by any amount. Under this definition, 166,865,198 bp of the naive per-sample union (86.12% of 193,759,589 passing bp) fell within intervals overlapping a cCRE. The recalibrated merged-max filter retained 84,814,444 bp in cCRE-overlapping intervals, representing 92.06% of its 92,129,994 passing bp. The merged FDR recalibration reduced the total called genomic space but increased the proportion of its retained intervals intersected ENCODE cCRE annotations. This enrichment is consistent with improved precision.

### Supplemental figures

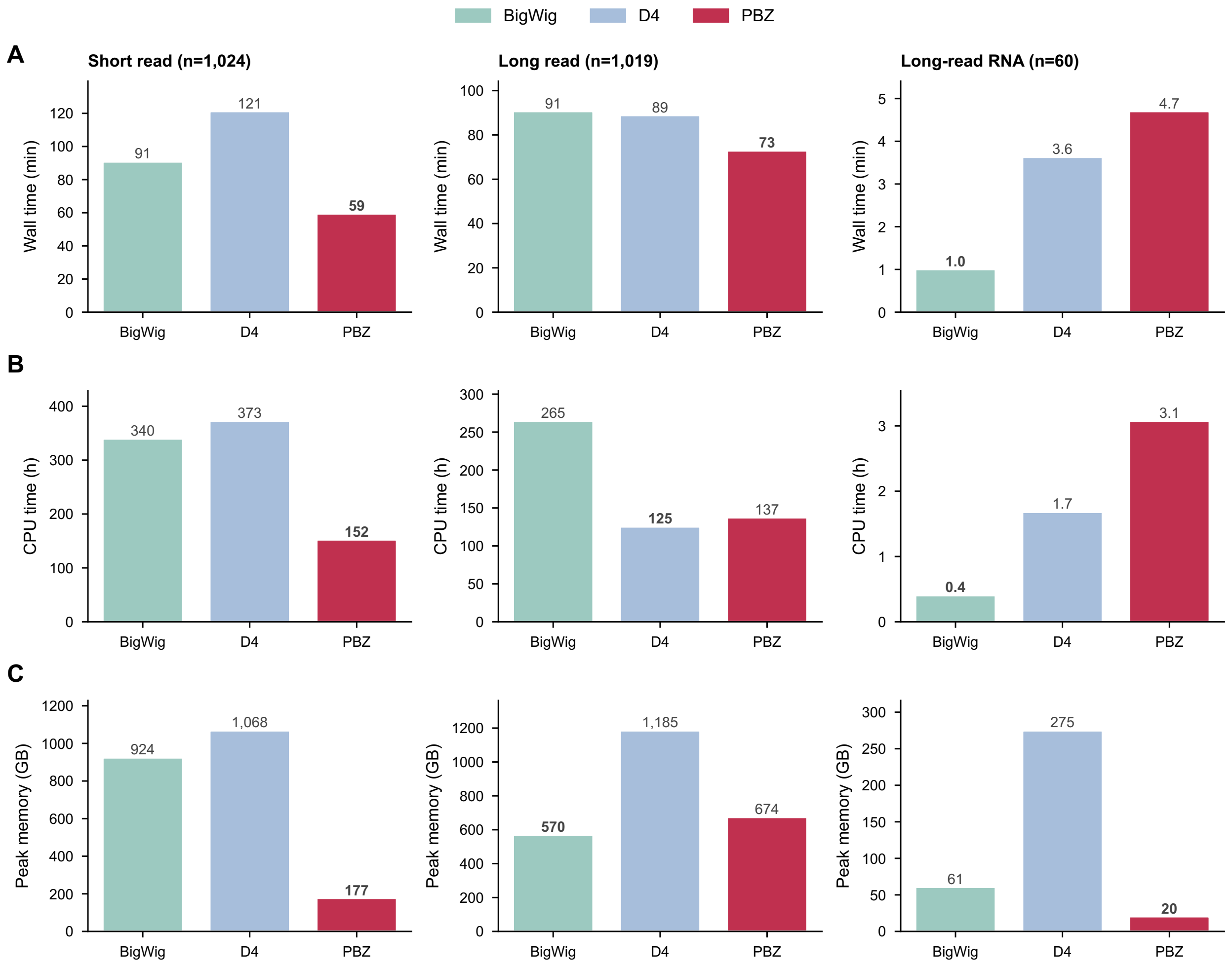


**Supplementary Figure S1.** Cost of building the largest cohort of each data type. (A) Wall-clock time, (B) total CPU time and (C) peak memory for building each format's dataset from 1,024 short-read genomes, 1,019 long-read genomes and 60 long-read RNA samples, on 256 cores. For D4 and bigWig, CPU time and memory are summed over the per-sample jobs running at the same time. The lowest value in each panel is in bold.

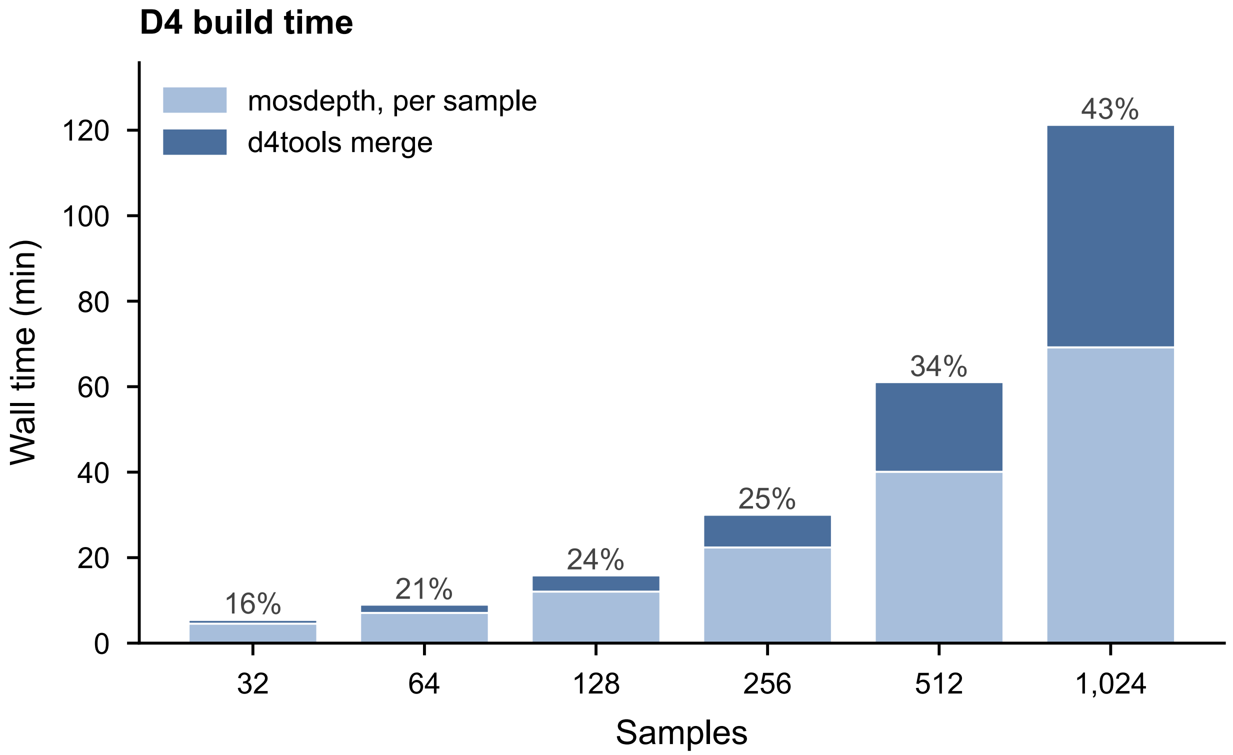


**Supplementary Figure S2.** Time spent merging per-sample D4 files. Wall-clock time to build the short-read D4 dataset at each cohort size, split into computing per-sample depth with mosdepth (light) and merging the per-sample files into one cohort file with d4tools merge (dark). Labels give the share of the build spent merging. Values are medians of repeated builds.

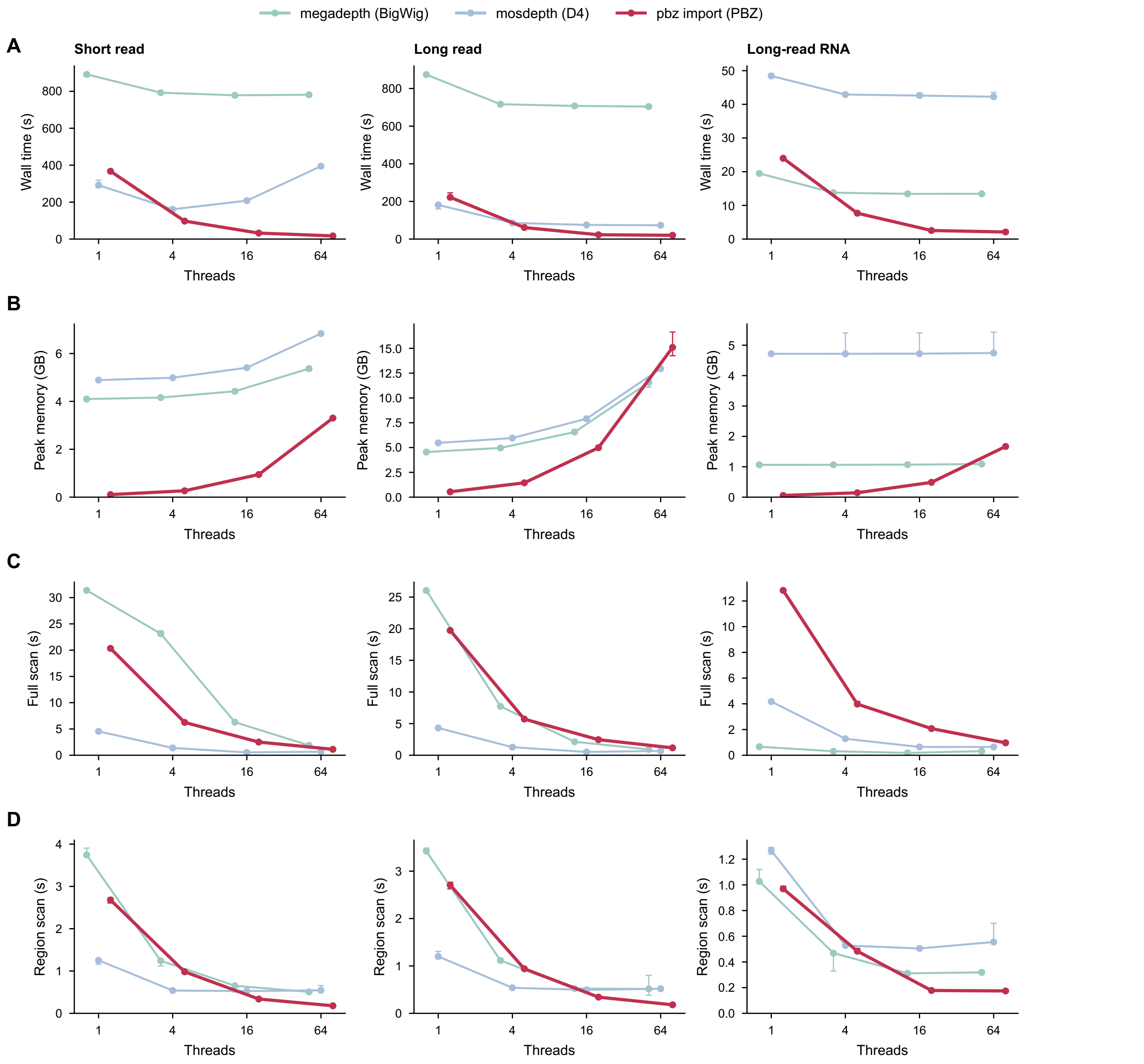


**Supplementary Figure S3.** Single-sample build and query performance. (A) Wall-clock time and (B) peak memory to compute per-base depth for one sample with megadepth (bigWig), mosdepth (D4) and pbz import (pbzarr), at 1, 4, 16 and 64 threads. Wall-clock time to (C) compute the mean depth over the whole genome (full scan) and (D) compute the mean depth over 10,000 random 10-kb regions (region scan), on each tool's output. The pbzarr and D4 scans used 1, 4, 16 and 64 threads, and the bigWig scan used one thread. For each data type, we used the sample whose input file was the median size: among the first 1,024 short-read samples, HG00475 (15.9 GB CRAM); among all 1,019 long-read samples, NA18539 (5.5 GB CRAM); and among all 60 long-read RNA samples, HG00260 (2.2 GB BAM). Values are medians of three runs (A, B) or three trials (C, D), each starting with the input evicted from the page cache.
